# Transcriptome-wide association study and fly experiments uncover the role of *CoA synthase* in ageing across species

**DOI:** 10.64898/2026.02.02.703301

**Authors:** Georgina Navoly, Aisyah Alizan, Olga Giannakopoulou, Stefanie Mueller, Linda Partridge, Nazif Alic, Karoline Kuchenbaecker

## Abstract

While significant progress has been made in understanding the genetic architecture of ageing in model organisms, our understanding of human ageing remains limited. We performed a multi-tissue Transcriptome-wide association study (TWAS) on human lifespan, integrating GWAS data from >1 million parental lifespans with gene expression prediction models derived from reference transcriptomic datasets; followed by replication using healthspan and longevity phenotypes as additional readouts of ageing. The TWAS uncovered 563 significant gene associations, of which 139 replicated. *TOMM40*, encoding a component of the mitochondrial outer membrane translocase that is fundamental for mitochondrial function, had the strongest association with parental lifespan and longevity and was fine-mapped as a putatively causal lifespan gene at the *APOE-TOMM40* region. Uniquely in our study, we identified fly orthologues of replicating genes and examined if modulating their expression impacts *Drosophila* longevity. The nine novel associations with all three ageing outcomes included *COASY,* encoding Coenzyme A synthase. Knocking down its fly orthologue, *Ppat-dpck*, resulted in significant lifespan extension in flies. Hence, in addition to discovering new genes associated with human ageing, by combining human TWAS with experimental *Drosophila* work, we provide evidence for the role of *COASY* (*Ppat-dpck*) in ageing across species.

**Significance statement:** Extensive research on ageing has been conducted in *Drosophila* and *C.elegans* due to their short lifespan and experimental tractability. However, in human genetic research, only a few loci have been consistently replicated. To bridge this gap, we conducted a transcriptome-wide association study (TWAS) followed by experimental validation in *Drosophila*. TWAS revealed 139 significant and replicating gene associations, including *COASY.* Knockdown of its fly orthologue (*Ppat-dpck*) significantly extended fly lifespan, validating its roles in ageing across species. Thus, integrating multiple ageing outcomes through TWAS in human genetic research can uncover robust associations and highlight genes involved in fundamental ageing mechanisms.

## Background

Most but not all animals age^1^. Ageing is thought of as an intrinsic process which manifests in a number of ways: decline in physiological functions, increased susceptibility to disease and increased probability of death with time. Extensive research on ageing has been conducted using animal models, such as *Drosophila melanogaster* and *Caenorhabditis elegans,* due to their short lifespan and experimental tractability. These research efforts have demonstrated that the rate of ageing can be modified^2–6^. Indeed, results of animal studies have revealed the involvement of several biological pathways in the ageing process^7–11^. While we have a substantial understanding of the genes and processes contributing to ageing in model organisms, our understanding of the genetics and biology of human ageing is lagging behind.

Genome-wide association studies (GWAS) have extensively been used to understand the genetic basis of complex traits in humans, including ageing^12^. Previous GWAS have identified multiple genetic variants associated with human lifespan, however, only a few genome-wide significant loci affecting lifespan have been repeatedly highlighted and replicated across studies, including *APOE, CDKN2A/B, LDLR, LPA, CHRNA3/5*^13–20^. These genes are involved in cellular senescence, lipid metabolism, immunity and inflammation pathways and are associated with age-related diseases, such as cardiovascular disease^21–24^. Although several genes have been identified that play a role in ageing in model organisms, and a few genes that play a role in human ageing, there is little documented overlap between the two. *APOE*, for example, the most robustly replicated ageing-related gene in humans, does not have direct orthologues in invertebrate models, and its study has even been complicated in mice due to species differences^25,26^.

As a new attempt to combine work in humans and model organisms, we performed a transcriptome-wide association study (TWAS), followed by experimental manipulation of select genes in *Drosophila*. TWAS is a gene-based association analysis that tests whether inherited differences in gene expression are linked to an outcome. By leveraging the rich biological information contained in gene expression data, TWAS can be particularly effective in providing better insights into the genetic basis of complex traits that are not well explained by GWAS alone^27–29^. Additionally, as a large proportion of SNPs identified in GWAS are likely regulatory in nature, additional studies are required to determine the specific genes they influence^12^; TWAS can tackle this question directly.

## Results

### Study design

We performed multi-tissue TWAS using three different ageing outcomes, with gene expression data from 48 human tissues from the Genotype-Tissue Expression project (GTEx)^30^. We used parental lifespan as the ageing outcome for our discovery study, leveraging GWAS data from over 1 million parental lifespans from the UK Biobank (UKB) and the LifeGen Consortium. We then performed two validation TWAS analyses: using healthspan (morbidity-free lifespan), with data from the UKB; and for the external replication TWAS (data not from UKB), we used longevity as the ageing outcome (central section of Figure 1). We used the TWAS fine-mapping tool FOGS, to fine-map our genes and prioritise the TWAS associations (left-hand side of Figure 1). We then directly tested the role in ageing of the fly orthologues of four genes associated with human ageing in the TWAS to assess whether these genes are implicated in ageing across species (right-hand side of Figure 1).

### TWAS of parental lifespan

**Figure 1.**
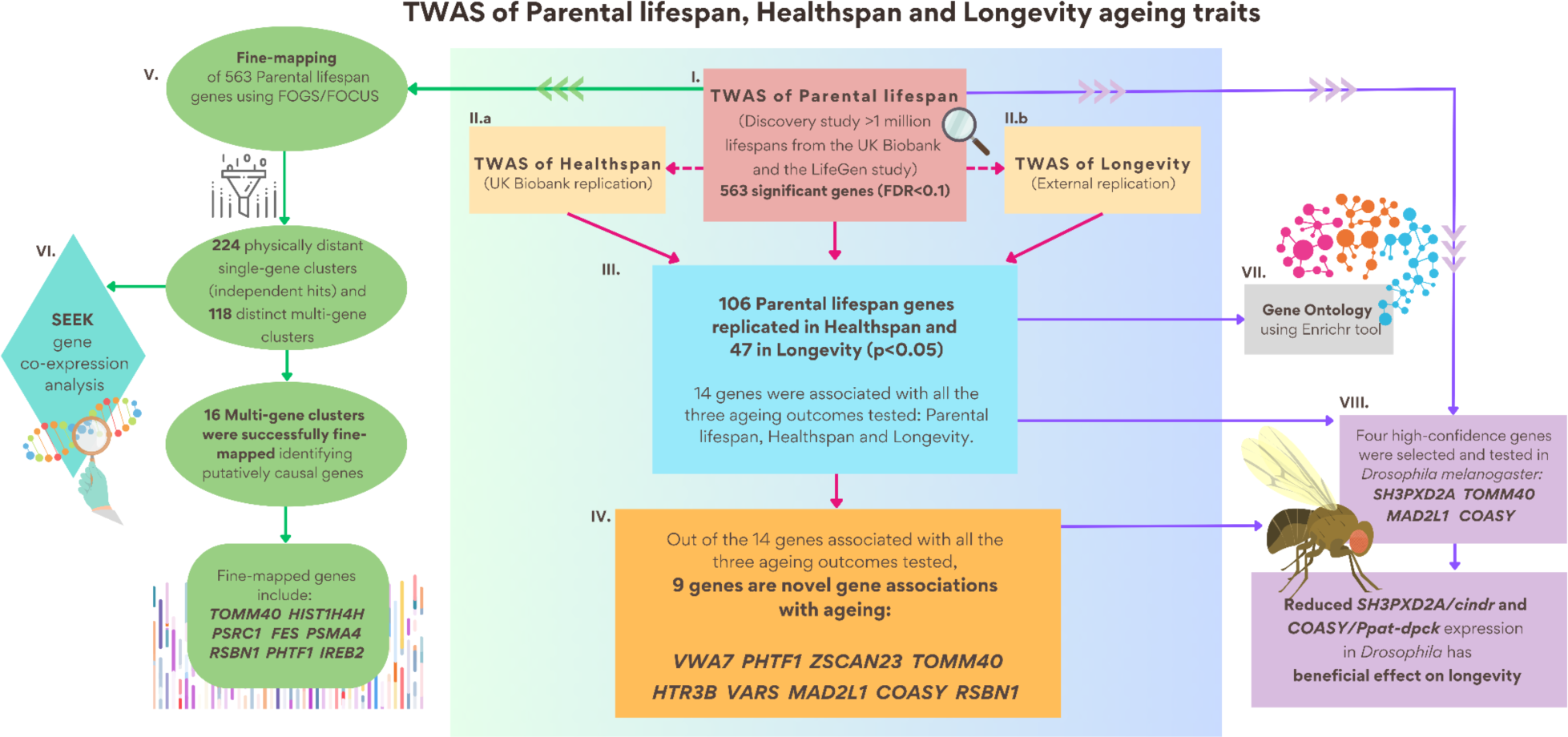
Summary of analyses. The central panel shows the TWAS results. The multi-tissue discovery TWAS identified 563 significant genes (False discovery rate (FDR) <0.1) associated with parental lifespan (middle panel, in red). In the healthspan TWAS, 214 genes were significantly associated with healthspan (FDR < 0.1), while in the third multi-tissue ageing TWAS, 7 genes showed significant associations with longevity (FDR < 0.1) (middle panels, in yellow). These TWAS results were not considered on their own, but were used as replication studies to replicate findings from the parental lifespan discovery TWAS. 139 parental lifespan genes were replicating in at least one other ageing outcome (106 replicated in healthspan and 47 in longevity, p<0.05) (middle panel, in blue). Replicating genes were used for Gene Ontology analyses, and their role in lifespan modulation were investigated experimentally in *Drosophila melanogaster* (right-hand side panels). The 563 genes associated with parental lifespan were located in 118 physically distinct (+/-100kb) multi-gene clusters. FOGS TWAS fine-mapping tool was used to fine-map these clusters (left-hand side panel).

The multi-tissue discovery TWAS identified 563 significant genes (False discovery rate (FDR) <0.1) associated with parental lifespan (Figure 2, Supplementary Table 1). This included genes in close proximity to all the 12 genome-wide significant single-variant loci from the original GWAS^31^ (*APOE* (p=9.68E-60), *CHRNA3* (p=4.11E-26)/*CHRNA5* (p=2.57E-23), *LPA* (p=1.50E-23), *FES* (p=2.04E-09)/*FURIN* (p=3.43E-06), *LDLR* (p=3.65E-07), *HTT* (p=4.34E-07), *HLA-DQA1* (p=1.07E-06), *HP* (p=2.09E-06), *KCNK3* (p=3.25E-06), *ATXN2* (p=1.54E-05), *MAGI3* (p=3.83E-05), *CDKN2B* (p=4.32E-05)). We also identified 109 novel genes that have not been previously implicated in human ageing, including *TOMM40* (p=2.79E-66), *HIST1H4H* (p=6.11E-08) and *SLC22A7* (p=1.73E-08) (Figure 2, Supplementary Table 1).

**Figure 2.**
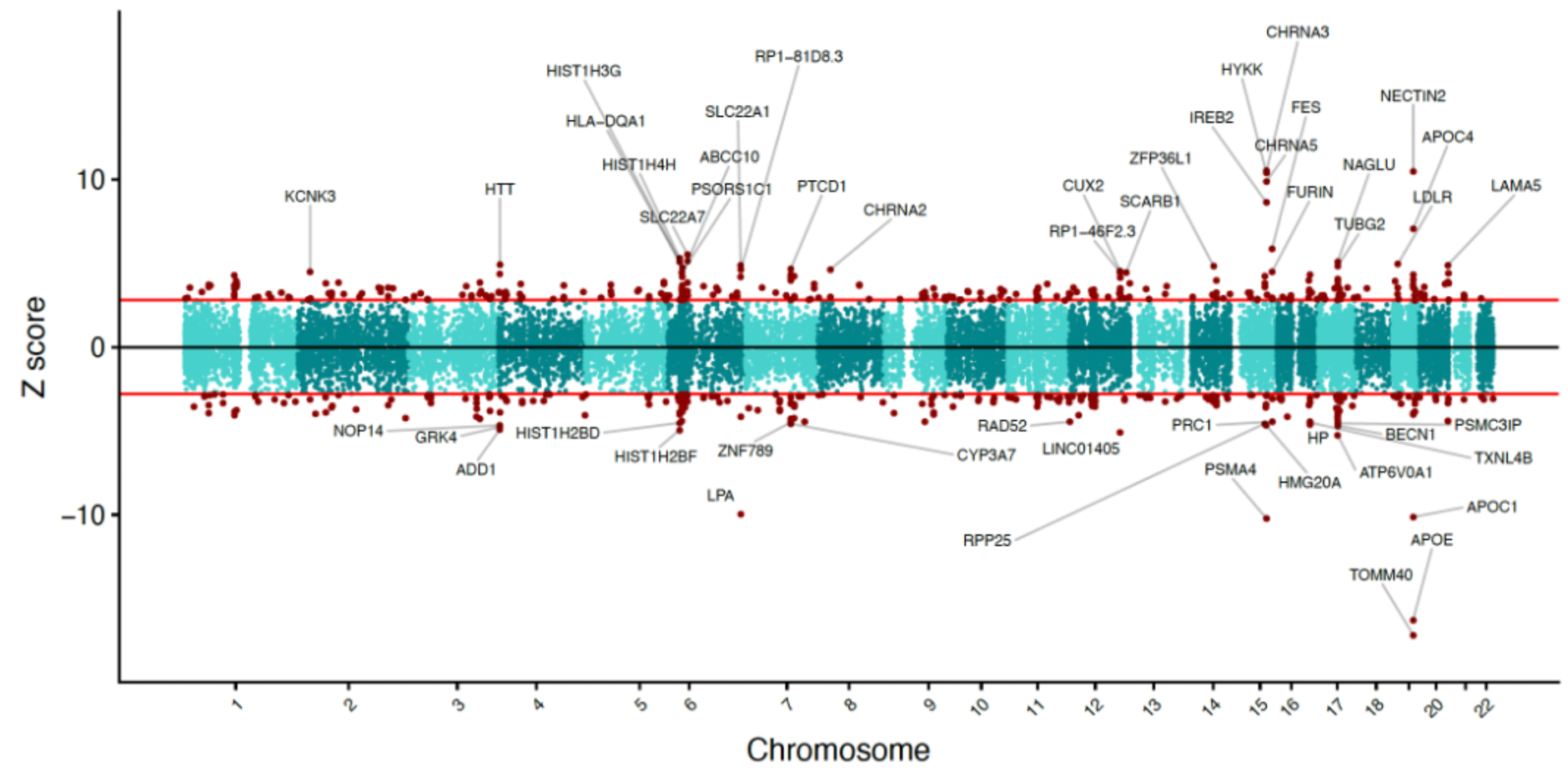
Manhattan-style z-score plot showing 22,316 gene associations (green and red dots), including the 563 statistically significant genes (red dots) associated with parental lifespan. The 50 most significant genes are annotated on this plot. The horizontal red lines mark the z-score significance threshold (consistent with FDR= 0.1 threshold of this dataset). Genes on the top part of the graph have a positive direction of effect (higher expression is associated with longer parental lifespan), while the genes on the bottom part of the plot have a negative direction of effect (inverse effect - higher expression is associated with shorter parental lifespan). Some of the 14 genes that show significant associations across all ageing outcomes tested are not displayed here, as they do not rank among the top 50 in the parental lifespan dataset.

### TWAS fine-mapping successfully resolves several key ageing-associated loci, proposing putative causal genes

Of the 563 genes associated with parental lifespan, 224 were located in physically distant loci with only one gene within +/- 100kb, including *SH3PXD2A, ZSCAN23, MAD2L1 and HTR3B.* These genes were considered independent hits. The remaining 339 genes were located in 118 distinct (< +/- 100kb) multi-gene clusters. As TWAS may identify multiple gene-trait associations in a region due to linkage disequilibrium among single nucleotide polymorphisms, we used the FOGS to fine-map these multi-gene clusters and prioritise the most likely causal genes. We successfully resolved 16 loci to one high-confidence target gene each, including *HIST1H4H, FES, PSRC1* (Supplementary Table 2). To consider a fine-mapping result valid, the top two genes in each multi-gene cluster, with the smallest p-values, had to be available from the same tissue(s). In gene clusters where this condition was not met, we concluded that the genomic region cannot be resolved. While the inclusion of this stringent quality control step was important to produce reliable fine-mapping results, it also reduced our ability to resolve a higher number of loci.

The *APOE-*region has previously been shown to be important in ageing^13^. In our work, in the multi-gene cluster at 19q13 *(APOE, TOMM40, NECTIN2, APOC1, BCAM)*, *TOMM40,* a gene nearby *APOE* was prioritised as the putatively causal lifespan gene (Posterior inclusion probability (PIP))= 0.99. *APOE* also yielded a posterior probability of 20%, while the rest of the genes in the cluster had PIP score of <0.00016 (0.0016% to be putatively causal) (Supplementary Table 2).

As a sensitivity analysis, we performed SEEK co-expression analysis of the genes classified independent (based on their location) to confirm that these genes are not co-expressed (Supplementary Table 3). 220 out of the 224 genes tested were not co-expressed with their neighbouring gene(s) and were therefore confirmed as independent by SEEK. Four genes were indicated to be significantly co-expressed by SEEK (SEEK co-expression score (SCES) > 1.96): *CRYGC* was co-expressed with gene *GBX2* from cluster immediately next to it (SEEK co-expression score (SCES)= 2.175), *PHIP* was co-expressed with *PNISR* (SCES= 2.69), *PNISR* was co-expressed with *PHIP* (SCES= 2.75) and *CENPP* was co-expressed with *SMC2* from the cluster immediately next to it (SCES= 2.36 (Supplementary Table 3)).

### Replication across ageing outcomes

Since ageing can manifest in myriad ways in humans, different types of ageing outcomes have been studied. We hypothesised that genes that are linked to fundamental ageing mechanisms and influence core components of ageing, would be associated across different ageing outcomes; our work is the first to include multiple ageing outcomes in a TWAS. In the healthspan TWAS, 214 genes were significantly associated with healthspan (FDR < 0.1), while in the third multi-tissue ageing TWAS, 7 genes showed significant associations with longevity (FDR < 0.1) (Supplementary Tables 4 and 5). These TWAS results were not considered on their own but were used as replication studies to replicate findings from the parental lifespan discovery TWAS, where the replication p-value of 0.05 was considered significant. We assessed the replication of the 563 genes significantly associated with parental lifespan in TWAS of healthspan (n=300,447; UKB replication) and longevity (n=11,262; non-UKB replication) as ageing outcomes (Supplementary Tables 4 and 5). We identified 139 significant (replication threshold p<0.05), replicating gene associations which were linked with parental lifespan and either healthspan or longevity (Table 1, Supplementary Tables 6 and 7). Out of these, 109 novel gene associations have not been implicated in previous GWA studies. 84 of these replicated in the healthspan TWAS and 34 in longevity, likely due to the considerably smaller sample size of the latter. 14 genes (9 of which are novel gene hits) replicated in both the healthspan and longevity outcomes (Table 1, top panel).

**Table 1.**
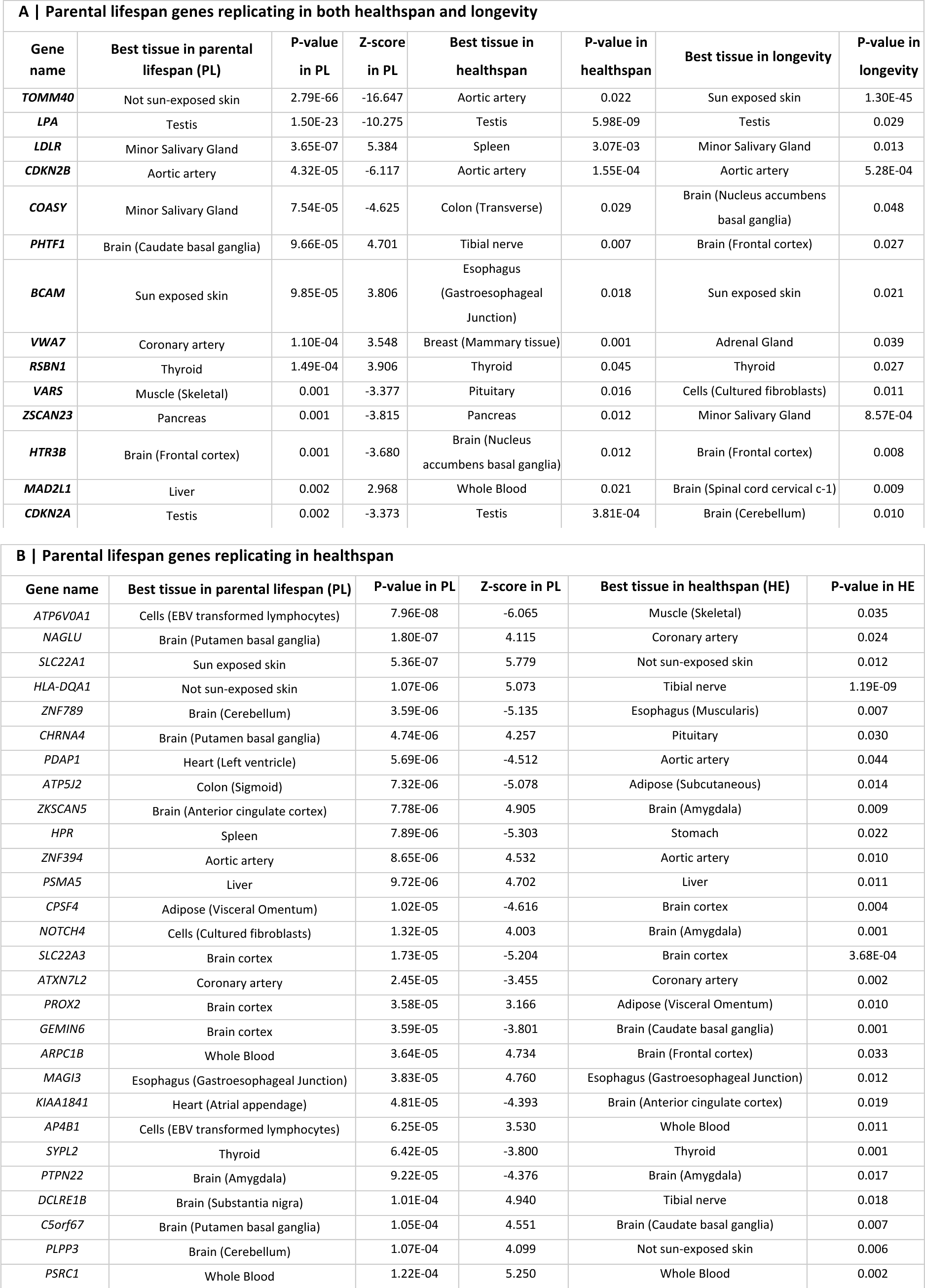
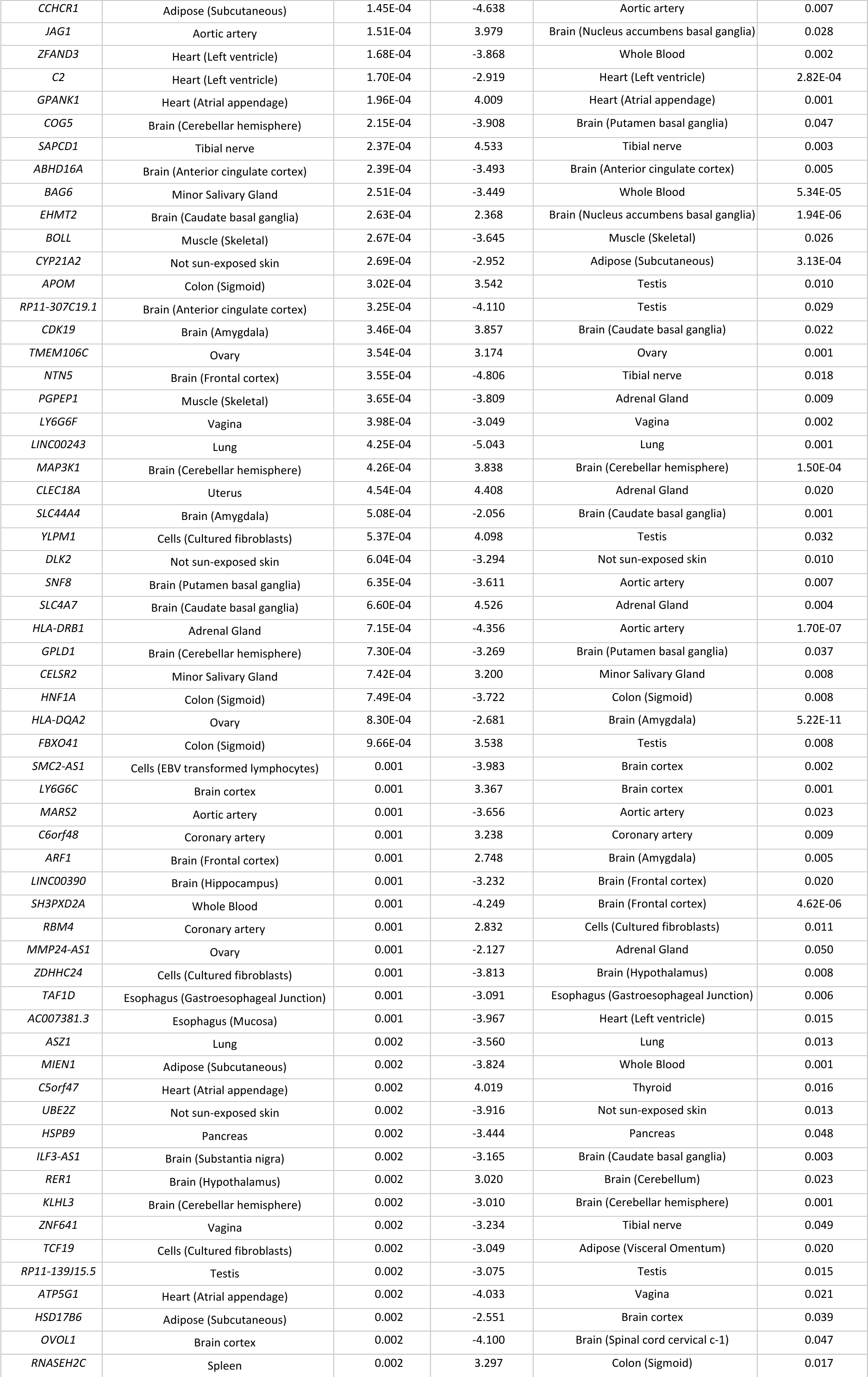
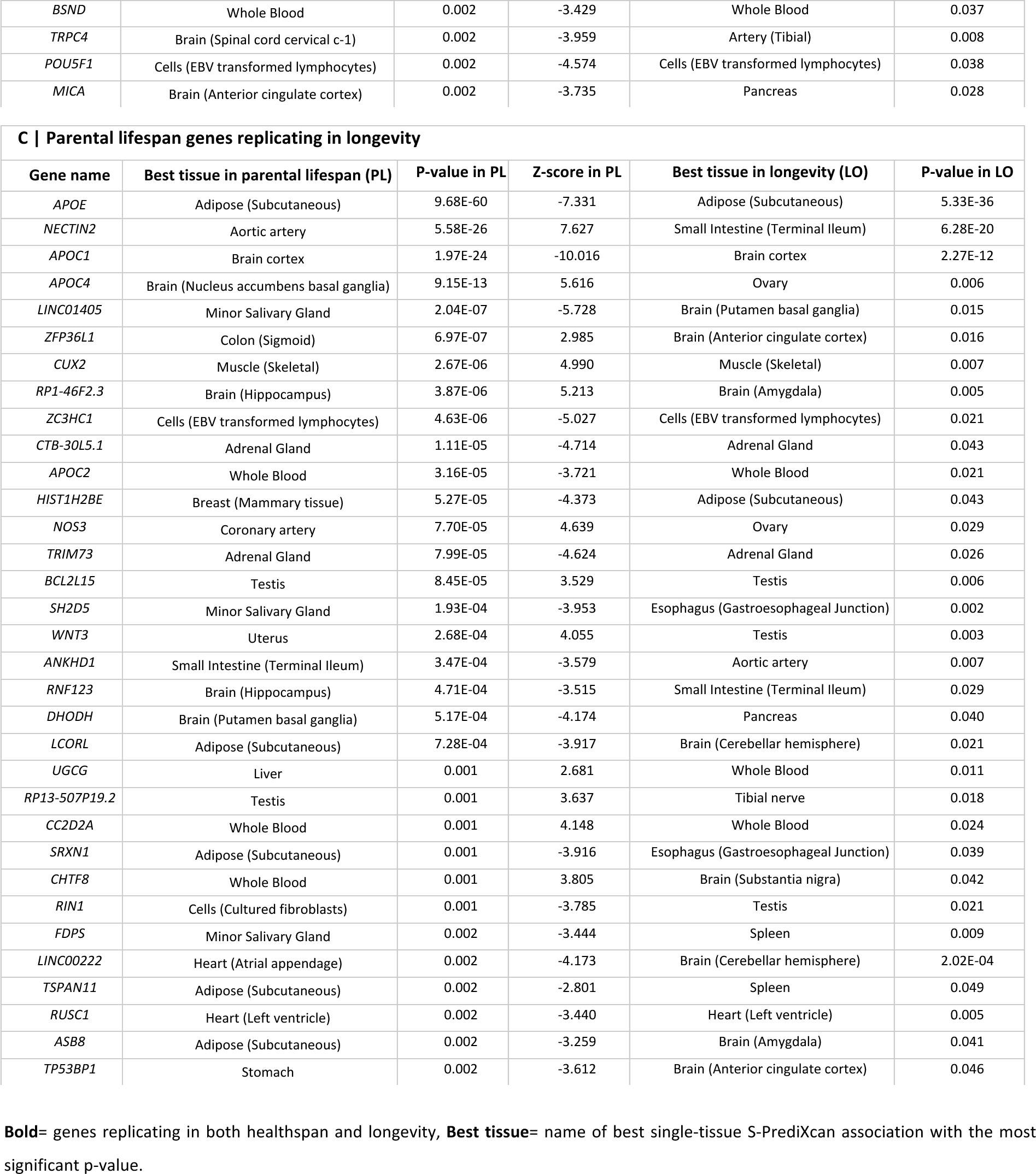
Replicating genes associated in the TWAS of parental lifespan, showing 14 genes associated with parental lifespan, healthspan and longevity (top panel) and 139 genes associated with at least two ageing outcomes.

The nine novel gene associations with parental lifespan, replicating in both healthspan and longevity, are involved in biological functions important in ageing, including cognition and memory (*HTR3B*), maintaining spindle checkpoint function (*MAD2L1*) and playing an important role in synthetic and degradative metabolic pathways (*COASY*). *TOMM40*, a gene that has an important role in maintaining mitochondrial function, displayed the strongest gene association with parental lifespan (p=2.79E-66, z-score (z)= -16.64, best tissue: Not sun exposed suprapubic skin), as well as with longevity (p=1.30E-45, z= -14.42, best tissue: Sun exposed lower leg skin) at the *APOE-TOMM40* region, showing more significant associations in terms of its expression than *APOE* (p=9.68E-60, z= -7.33 and p=5.33E-36, z= -6.75, respectively). *TOMM40* also significantly replicated in healthspan (p=0.022, at a replication threshold of 0.05), while *APOE* did not show significant replication in healthspan. Another gene association with all three ageing outcomes is *HTR3B* (p*=*0.0013 / z= -3.68 in parental lifespan, best tissue: Brain Frontal Cortex; p=0.012 in healthspan replication, best tissue: Brain Nucleus accumbens basal ganglia; and p=0.008 in longevity replication, best tissue: Brain Frontal Cortex) (Table 1). Gene *HTR3B* encodes a subunit of 5-HT3 receptor (serotonin), a key neurotransmitter and an important target for the most commonly prescribed group of antidepressants.

### Gene Ontology analyses of ageing associated genes

To annotate biological mechanisms of parental lifespan genes replicating in healthspan and longevity ageing outcomes, we performed a Gene Ontology (GO) analysis (Figure 3, Supplementary Tables 8 and 9).

**Figure 3.**
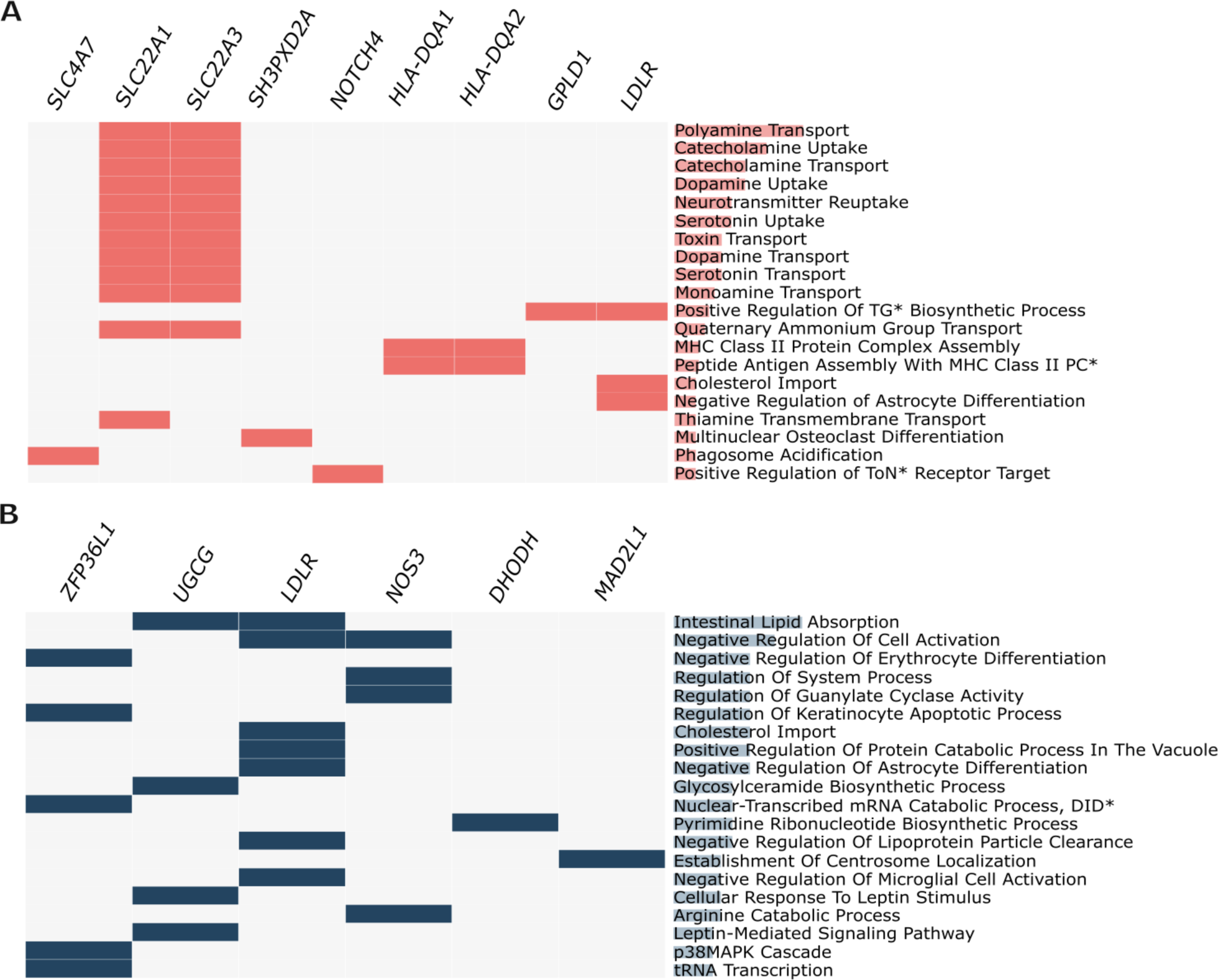
Gene Ontology analyses of parental lifespan genes replicating in healthspan and longevity. A) Bar plot showing the top 20 enriched GO terms for parental lifespan genes replicating in healthspan. **B)** Bar plot showing the top 20 enriched GO terms for parental lifespan genes replicating in longevity. The length of the horizontal bars covering the GO categories represent the Combined Score, which is a key ranking metric used by Enrichr. Combined Score = log(p-value) × z-score, where the z-score is based on the deviation from the expected rank - longer bars indicate stronger enrichment^32^.

In the healthspan replication analysis, *SLC22A1 and SLC22A3* showed the strongest associations from Gene Ontology for a number of GO terms, including Polyamine transport, Dopamine uptake, Serotonin uptake and transport (Figure 3A, Supplementary Table 8). The longevity replication results were significantly enriched in GO terms including lipid metabolism and immune response related pathways (Figure 3B, Supplementary Table 9). Our GO analysis highlights key pathways linked to immune function that are consistent with previous findings, strengthening the evidence that age-related changes in immune function are a strong feature of ageing^33^. The full GO results for parental lifespan genes replicating in healthspan and longevity are available in Supplementary Tables 8 and 9.

### Knock-down of *Ppat-dpck (COASY in humans)* in fly neurons extends lifespan

We sought to experimentally query the role in animal ageing for a selection of the human genes using the fruit fly as a discovery platform. The fruit fly is a powerful genetic model where gene expression can be experimentally altered in specific cell types of the adult using the GeneSwitch system^34–36^ combined with either RNA interference (RNAi) or overexpression constructs to reduce or increase gene expression, providing an experimental system that can mimic TWAS findings. We chose four high-confidence and biologically relevant genes: *COASY, MAD2L1, TOMM40, SH3PXD2A* based on both our findings and practical consideration: the four were associated with at least two ageing outcomes and they each had a fly orthologue, namely *Ppat-dpck, Mad2, Tom40* and *cindr*, for which suitable reagents were available. The fly genes showed a variable degree of homology to the human counterpart (Supplementary Table 10ab). For the genes where the TWAS analysis indicated that reduced expression is associated with slower ageing, we designed experiments to knockdown the fly orthologue (*cindr, Ppat-dpck,* and *Tom40*). On the other hand, an overexpression line for *mad2* was used as increased expression was predicted by TWAS to be beneficial. The genetic reagents available for each gene were validated by qPCR (Supplementary Table 11, Supplementary Figure 1). The specific tissues targeted were chosen based on which human tissues gave strong signals for association to ageing in the TWAS, and practical consideration of the tools (drivers) available in flies which are limited to certain cell types, tissues or organs. Human brain tissue had a strong association to ageing outcomes for *COASY, MAD2L1* and *SH3PXD2A*; the neurons were chosen as the relevant cell type in *Drosophila*. The transverse colon tissue was also associated with ageing for *COASY*, hence, we also knocked down its orthologue *Ppat-dpck* in the fly gut. For *TOMM40*, the implicated best tissues were the skin and aorta. Due to lack of more specific drivers, we chose to knockdown *Tom40* ubiquitously. Note that all four orthologues show similar expression levels across different cell types in both flies and humans, with no substantial evidence of cell-type restricted expression (Supplementary Figure 2 and 3). To avoid effects on development and to mimic more closely the findings of TWAS, which is based on adult expression data, we performed all manipulations starting from day two of adulthood.

We chose to examine lifespan as this is the canonical readout of ageing in experimental ageing studies. We specifically focused on females as their lifespan appears to be more plastic (modifiable) in the fruit fly^37,38^. Indeed, interventions that have been discovered in fruit fly females work in both sexes of mice^4,10,38–41^. Knocking down *Ppat-dpck* (best orthologue of *COASY*) and *cindr* (best orthologue of *SH3PXD2A*) in fly neurons was sufficient to cause a significant extension in lifespan (p=0.004 and p=6x10^-16^, respectively, Figure 4 A-C, Supplementary Table 12)). Feeding of the inducer (RU486) in the absence of the RNAi constructs did not extend lifespan, confirming that the longevity is not an artefact of RU486 feeding (Figure 4A, C). When we knocked down *Ppat-dpck* specifically in the fly gut, we saw no difference in lifespan (p=0.121, Figure 4D), indicating the effects of *Ppat-dpck* on longevity are tissue specific. We found no evidence for the role of *Tom40* (best orthologue of *TOMM40*) or *mad2* (best orthologue of *MAD2L1*) in fruit fly longevity (Supplementary Figure 4). Note that these negative results do not necessarily mean that *Tom40* and *mad2* are not implicated in fruit fly longevity. It is plausible that a different level of knockdown or in a specific or different tissue would have revealed an effect, as further studies may show.

Note that *cindr* does not share high homology with *SH3PXD2A* (19% amino acid identity, Supplementary Table 10b) and the homology is limited to a portion of the protein encoded by *SH3PXD2A* (Figure 4B), making it likely that *cindr* only recapitulates a part of *SH3PXD2A* function; still, our experiments indicate that this shared function may be relevant for animal ageing. On the other hand, *Ppat-dpck* is a clear, direct orthologue of the human *COASY* (37% amino acid identity; Supplementary Table 10b) and they both encode the last enzyme in the coenzyme A biosynthesis pathway^42^. Furthermore, flies mutated in *Ppat-dpck* display phenotypes consistent with the role of this gene in CoA biosynthesis ^43^. We additionally examined whether the role of *cinder* and *Ppat-dpck* in fruit fly ageing can be replicated in a different genetic background. Our initial screen was performed in a hybrid genetic background, resulting from crossing the driver which was carried in an outbred, healthy wild-type background, with the responder lines which were generated in an inbred strain. We backcrossed *UAS-cindr^RNAi^ UAS-Ppat-dpck^RNAi^* six times into the outbred background and repeated the lifespan assay, to ensure the longevity can also be observed in a natural, healthy, outbred background and is not simply rescuing any artificial, inbreeding effects. While knockdown of *cindr* still showed an extension of median lifespan, the overall difference in survival was diminished and failed to pass the formal statistical significance threshold of p<0.05 by a small margin (p=0.063, Figure 4E), indicating that neuronal *cindr* knockdown has a smaller effect in the outbred background. On the other hand, we found that the knockdown of *Ppat-dpck* in adult neurons could extend lifespan even in this healthy, outbred genetic background (Figure 4F), indicating that *Ppat-dpck* has a robust effect on longevity. In both experiments, the survival of the driver alone control was not impacted by RU486 feeding (Supplementary Figure 4C and D). Overall, two out of four genes tested could impact longevity in the manner consistent with the TWAS predictions This provides validation for our TWAS as the proportion (50%) is substantially higher than would be expected by chance: unbiased genetic screens in *C. elegans* indicate that the proportion of longevity genes is approximately between 0.1-2.4%^44,45^, and this is similar to the number of genes annotated as determining adult lifespan in *Drosophila* (1.5%) and *C. elegans* (2.3%).

**Figure 4.**
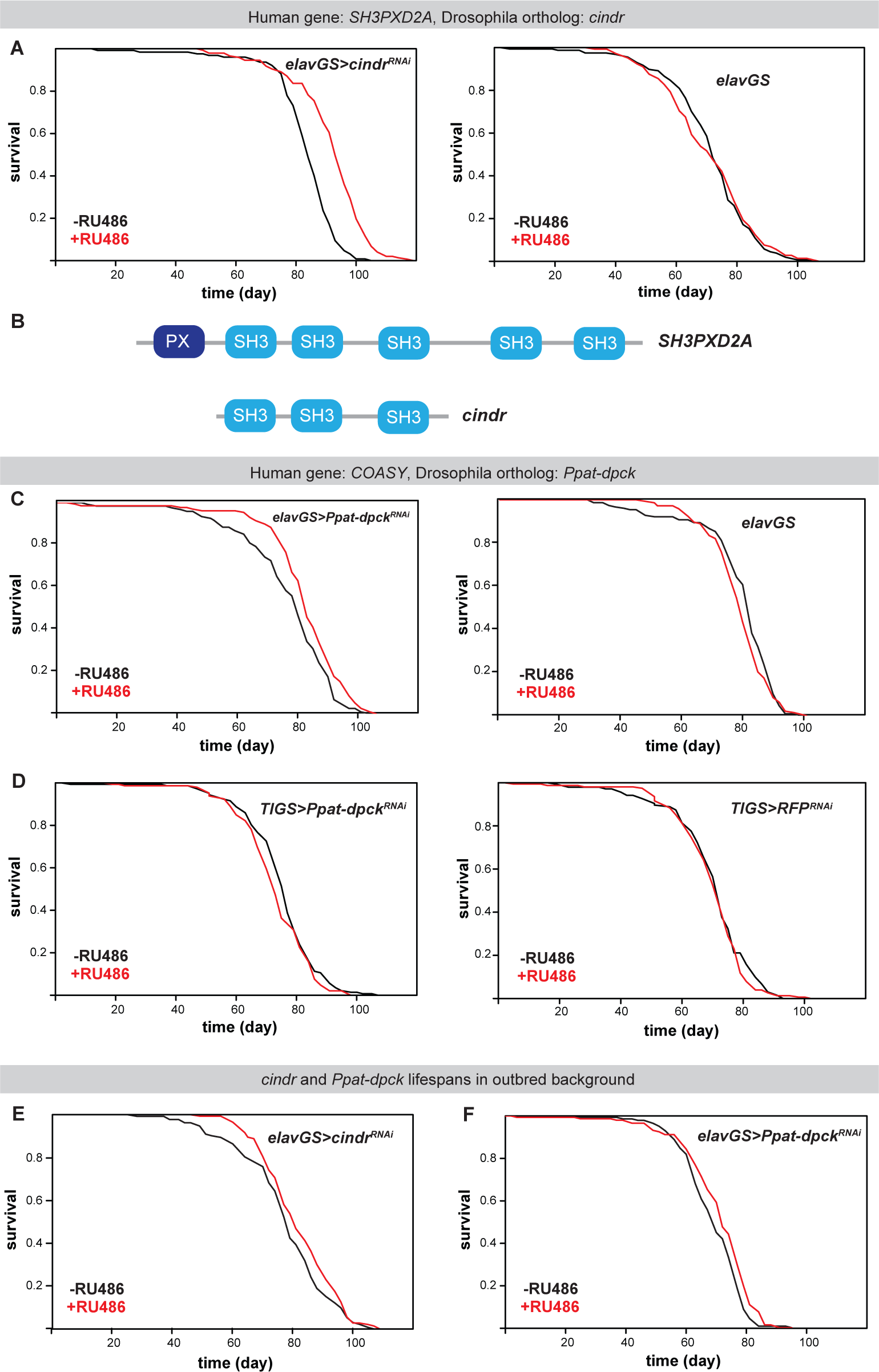
Inhibition of *cindr* in neurons extends lifespan. **(A)** Lifespans of female flies with *cindr^RNAi^* induced in neurons by feeding RU486 (-RU486 n = 127 dead/0 censored flies, median lifespan = 84 days, last quantile = 89 days, +RU486 n = 147 dead/4 censored flies, median lifespan = 93 days, last quantile = 100 days, p = 6.1 x 10^-16^, log-rank test) and the wild-type controls (-RU486 n = 157 dead/0 censored flies, median lifespan = 72 days, last quantile = 82 days, +RU486 n = 145 dead/0 censored flies, median lifespan = 72 days, last quantile = 82 days, p = 0.893, log-rank test). **(B)** Domain organisation of the human *SH3PXD2A* and *Drosophila* cindr proteins. **(C)** Lifespan of female flies with *Ppat-dpck^RNAi^* induced in neurons by feeding RU486 (-RU486 n = 120 dead/4 censored flies, median lifespan = 80 days, last quantile = 87 days, +RU486 n = 125 dead/6 censored flies, median lifespan = 83 days, last quantile = 90 days, p = 0.004, log-rank test) and the wild-type controls (-RU486 n = 143 dead/11 censored flies, median lifespan = 83 days, last quantile = 87 days, +RU486 n = 144 dead/1 censored flies, median lifespan = 80 days, last quantile = 85 days, p = 0.133, log-rank test). **(D)** Lifespan of female flies with *Ppat-dpck^RNAi^* induced in the gut by feeding RU486 (-RU486 n = 142 dead/7 censored flies, median lifespan = 77 days, last quantile = 81, +RU486 n = 146 dead/0 censored flies, median lifespan = 73 days, last quantile = 81 days, p = 0.121, log-rank test) and the controls driving RFP expression (-RU486 n = 133 dead/10 censored flies, median lifespan = 73 days, last quantile = 77 days, +RU486 n = 153 dead/2 censored flies, median lifespan = 73 days, last quantile = 77 days, p = 0.384, log-rank test). **(E)** Lifespan of female flies with *Cindr^RNAi^* induced in the neurons in an outbred line by feeding RU486 (-RU486 n = 132 dead/0 censored flies, median lifespan = 79 days, last quantile = 86 days, +RU486 n = 143 dead/0 censored flies, median lifespan = 81 days, last quantile = 91 days, p = 0.063 **(F)** Lifespan of female flies with *Ppat-dpck^RNAi^* induced in the neurons in an outbred line by feeding RU486 (-RU486 n = 140 dead/5 censored flies, median lifespan = 70 days, last quantile = 77 days, +RU486 n = 147 dead/1 censored flies, median lifespan = 72 days, last quantile = 79 days, p = 6.91 x 10-3, log-rank test).

## Discussion

Our results demonstrated that TWAS can advance our understanding of the genetics of ageing by unveiling a large number of novel gene associations. Previous ageing GWAS have already included millions of human age-related outcomes. However, despite the large sample size, these studies only identified a few genes consistently associated with ageing. In our study, using existing human data, we introduced novel ways of studying ageing through TWAS in combination with experimental *Drosophila* work. Unlike GWAS, TWAS enabled the identification of a considerably larger number of genes associated with ageing and provided insights into tissues of interest and the direction of effects. This approach not only expanded our knowledge of ageing-related genes in humans, but it also facilitated direct testing of these genes in model organisms, such as *Drosophila* in this study. Notably, the genes identified through TWAS of parental lifespan, healthspan and longevity showed relatively distinct associations. The differences in genes we observed in each of the ageing outcomes might be due to the different properties of common diseases associated with the outcomes tested. Additionally, a recent multivariate analysis of multiple ageing traits (parental lifespan, healthspan and longevity) concluded that genetic loci that were not shared between traits often associated with population-specific, behavioural risk factors such as smoking^46^. As none of the outcomes fully capture the complex ways in which ageing can manifest, looking for genes replicating across different definitions may restrict findings to genes less likely linked to diseases or specific features of each outcome and more to fundamental ageing mechanisms. By including multiple ageing outcomes in our TWAS, our work extends previous research^47^, and improves our understanding of the genetics of ageing.

Remarkably, experimental manipulation of two of four genes tested was able to significantly extend lifespan in at least one genetic background in *Drosophila*, suggesting that combining multiple ageing outcomes may be a promising approach for future studies to target fundamental ageing genes, and emphasising the importance of forging closer connections between human genetic studies and experimental work in model organisms.

### *TOMM40*, a novel putative ageing-associated gene, independent of *APOE*

The *APOE* gene has been widely acknowledged as the most prominent ageing-related human gene association, consistently displaying the largest effect sizes, smallest p-values, and consistent replication across studies^13^. In our work, we showed that both *APOE* and the neighbouring gene, *TOMM40,* are associated with ageing. *TOMM40* encodes a protein in the outer membranes of mitochondria that is required for protein transport to the inside of the mitochondria^48^. Through *TOMM40’*s key role in maintaining mitochondrial function, it has been shown to influence age-related memory performance^49^. Healthy ageing is dependent on normal mitochondrial functioning. When mitochondria become dysfunctional, it can result in compromised energy metabolism, increased oxidative stress, and elevated levels of damage-associated molecular patterns that are released from damaged or stressed mitochondria^50^. These factors collectively contribute to biological changes associated with the ageing process^51^. In fact, most age-related diseases, particularly neurodegenerative diseases, have been linked to abnormal mitochondrial function, mediated through consequent cellular damage, likely leading to the degeneration of neurons^50^. In the majority of previous work in humans, *TOMM40* was associated with Alzheimer’s disease and neurodegeneration^52–54^. A recent study in *C.elegans* showed that the inhibition of *timm-23* or *tomm-40* increases lifespan in this model organism^55^ and a recent review suggested that certain *TOMM40* variants might potentially be associated with healthy ageing and longevity in humans^51^. However, neither of these studies provided strong evidence for the role of *TOMM40* in human ageing. In our work, we first showed *TOMM40* exhibiting a stronger and more significant association in terms of expression levels with ageing than *APOE*. We also highlight *TOMM40* as the prioritised, putatively causal gene in the region, over *APOE.* Whilst our work uncovered genes where expression levels associate with ageing, *APOE* has been shown to impact ageing through different functional alleles^14,56^. Therefore, both genes may play an independent role in ageing and our fine-mapping results also support this. *TOMM40* yielded a posterior probability of 99%, while APOE yielded 20%, which is substantially higher than an expected PIP score (<0.05) for a putatively non-causal gene.

### Elevated expression of Serotonin receptor encoding gene *HTR3B* is associated with shorter lifespan in humans

One of our other robust gene associations with all three ageing outcomes tested, was *HTR3B*. It encodes a subunit of 5-HT3 receptor, which binds serotonin, a key neurotransmitter and an important target for a key group of antidepressants in humans^57^. Previous studies using animal models showed that certain type of drugs that are used as antidepressants in humans and block the neural signalling of serotonin, can extend lifespan in C. *elegans*^58,59^ through mimicking dietary restriction and through the modulation of the insulin/IGF-like and TOR nutrient-sensing pathways, mechanisms well known to extend lifespan across species^60,61^. In addition, a recent study in *Drosophila* has shown that serotonin receptor signalling can modulate ageing in response to nutrient choice and showed that flies that were offered a single food (as opposed to multiple choices) produced less serotonin and lived longer lives, suggesting that food choices (or food availability) can affect lifespan through serotonin signalling^62^. Our findings support the role of serotonin signalling in ageing in humans. We showed that increased expression of gene *HTR3B* is associated with shorter parental lifespan, shorter healthspan and reduced longevity, indicating that the reduction of serotonin signalling in brain regions, specifically the frontal cortex and nucleus accumbens according to our TWAS results, is associated with longer lifespan and healthspan.

### Knock-down of *Ppat-dpck (COASY)* in fly neurons extends lifespan

While TWAS can uncover genes putatively involved in human ageing, experimental work in model organisms is required to cement their causal role in animal ageing. Most experimental studies are conducted independently of findings in human epidemiology, and findings often wait for years to be validated in humans. Here, using tools in fly genetics we showed that the fly orthologue of *COASY*, *Ppat-dpck*, and of *SH3PXD2A*, *cindr*, affect fruit fly lifespan, immediately validating two of our candidate genes and indicating functional conservation. The effects of *Ppat-dpck* knockdown were robustly replicated in an outbred, healthy genetic background. *Ppat-dpck* is the direct sequence and functional orthologue of the human *COASY*.

Both genes encode the bifunctional enzyme that catalyses the last steps in the Coenzyme A (CoA) biosynthesis pathway^43,63^. CoA synthesis is essential for organismal viability and function and yet we found that reducing *Ppat-dpck* expression in adult neurons promotes longevity. Such relationships are often observed in the genetics of ageing and are thought to be the result of antagonistic pleiotropy where gene activity is selected for beneficial effects that manifest early in life and have a strong impact on reproductive fitness even if they carry substantial costs for later health^64,65^. Interestingly, a reduction in acetyl-CoA in fly neurons by knockdown of acetyl-CoA synthetase increased fly lifespan, likely through enhanced autophagy^66^. It is plausible that the effect of *Ppat-dpck* and *COASY* on ageing are mediated by a related mechanism.

### Limitations

1. Our study has limitations. Currently, our TWAS fine-mapping analysis is based on a selection of tissues thought to be important in ageing. However, our current knowledge of the key tissues relevant to ageing is limited. In the future, better guidance on key tissues could improve the success of fine-mapping and help identification of more causal genes.
2. The current study is limited by the lack of both ancestral and geographic diversity due to the use of existing human genetics studies. GWAS and TWAS performed in more diverse groups exposed to diverse environments might detect additional, novel gene-trait associations and reduce the risk of identifying associations linked to common ageing risk factors such as smoking and common age-related diseases.
3. Three out of the nine high-confidence genes, *VWA7, ZSCAN23* and *VARS*, are located in the Major Histocompatibility Complex region, which might result in reduced robustness in variant-to-gene mapping due to high LD and promiscuous expression quantitative trait loci. However, the genes carried forward for validation in fly are not affected by this.
4. Lastly, the imperfect or limited orthology, as well as the limited possibility to manipulate fly tissues in a manner precisely equivalent to that of humans might have affected our experimental fly work.

## Conclusions

Inclusion of multiple ageing outcomes in human genetic research can uncover more robust associations and highlight genes involved in fundamental ageing processes. Combining, for the first time, human TWAS of ageing with experimental *Drosophila* work, we uncover the role of *COASY (Ppat-dpck)* in ageing across species. Our findings also elucidate the association of *TOMM40* expression with human ageing, independent of *APOE*.

## Methods

### Data resources and study design

In this study we used three different ageing outcomes. Hallmark biological characteristics of ageing for example DNA damage, telomere attrition, disturbed protein homeostasis, mitochondrial dysfunction, increased oxidative stress are challenging to measure directly in genetic association studies, thus human ageing is often measured through proxy phenotypes, including parental lifespan, healthspan, longevity^13^.

We used data from a parental lifespan GWAS for discovery in this study, with 512,047 maternal and 500,196 paternal lifespans of European ancestry from UKB and the LifeGen Consortium, where parental lifespan was defined as joint parental survival^31^. Parental lifespan is often used in place of participants’ lifespan, because the completed lifespans of most genotyped participants are usually not available. Thus, to increase the sample size and incorporate as many completed lifespans and long-lived survivors as possible into the analysis, the authors performed a survival analysis to transform parental survival into a quantitative trait and conducted a GWAS between parental survival and subject genotype for mothers and fathers separately^31^. The GWAS used 1,012,240 lifespans of parents of more than 500,000 genotyped participants in the UKB and the LifeGen consortium (A consortium of 26 population cohorts of European ancestry, with UKB lives removed) and found 12 genome-wide significant associations with parental lifespan^31^. For replication, we used data from a healthspan and longevity GWAS. The healthspan GWAS had data from 300,447 individuals with European ancestry from the UKB^17^. Healthspan was defined as the age of onset of the first disease from a cluster of seven common morbidities, strongly associated with ageing. The GWAS identified 12 genome-wide significant loci associated with healthspan^17^. Lastly, we used data from a longevity GWAS with 11,262 people with European ancestry, which compared individuals living to an age above the 90th survival percentile based on cohort life tables with those of average lifespan^14^. This study was based on 18 contributing cohorts and there was no sample overlap between the longevity GWAS and the discovery study, i.e. the parental lifespan GWAS. The GWAS identified 11 genome-wide significant loci associated with longevity, near genes *APOE, CDKN2B* and *CDKN2A*^14^.

### GWAS summary statistics quality control

GWAS summary statistics were filtered by removing variants with imputation accuracy less than 0.7. Variants with effect allele frequency below 0.01 and over 0.99 were also removed. Summary statistics were processed and formatted using the munge_sumstats.py script from the LD Score Regression (LDSC) software suite^67^.

### TWAS

A Transcriptome-Wide Association Study (TWAS) is a statistical approach that integrates genome-wide association study (GWAS) data with gene expression prediction models to identify genes whose expression is associated with a phenotype of interest. In this study, TWAS was performed using S-PrediXcan and S-MultiXcan, software packages to infer the relationship between genetically predicted gene expression and ageing outcomes. S-PrediXcan applies a two-stage regression method: first, it trains predictive models on genotype and gene expression data (for example from GTEx) to generate weights^68^. In the second stage, these weights are integrated with GWAS summary-level data to perform transcriptome-wide association analysis of predicted gene expression with phenotypes of interest^69^. S-MultiXcan, which requires single-tissue TWAS results from S-PrediXcan, extends this approach by combining information across multiple tissues, leveraging shared expression quantitative trait loci (eQTLs) to improve gene discovery. S-MultiXcan integrates evidence from multiple panels using multivariate regression, which incorporates the correlation structure^69^. In this study, Multivariate Adaptive Shrinkage in R-based (MASHR) prediction models were used. MASHR models utilise fine-mapping information and enhance statistical power by borrowing eQTL information across tissues and therefore are able to provide improved effect sizes, improved prediction and improved expression weights^70^.

We performed TWAS using S-PrediXcan and S-MultiXcan software in R (v4.3.4)^71^, which have been shown to predict gene expression more accurately than other TWAS methods^68,69,72^. To perform single tissue TWAS, a prerequisite of multi-tissue TWAS, the S-PrediXcan^73^ software was used. We used 48 GTEx V8 MASHR-based tissue models which are parsimonious, biologically informed and exhibit the greatest power^74^. We included tissues where the number of samples with genotype data per tissue were above 100 samples (as of April 2021), and therefore, excluded Kidney cortex, Bladder, Cervix - endocervix, Cervix - ectocervix, Fallopian tube and Kidney medulla.

We independently performed three TWAS (parental lifespan, healthspan, longevity) using 48 GTEx tissues (V8). To perform multi-tissue TWAS, we used the S-MultiXcan software^69^ which combines information about the strength of association between predicted expression and the phenotype of interest^69^.

### Fine-mapping of TWAS results

TWAS may identify multiple gene-trait associations in a region due to linkage disequilibrium among single nucleotide polymorphisms (SNPs) in the prediction models used to create SNP weights as well as co-expression. We used the TWAS fine-mapping tool FOGS in R (v4.3.4), to prioritise our gene hits and control for Type I error rates^75^. FOGS software computes a FOGS-aSPU (an adaptive test) score and also a FOCUS-PIP score based on a Bayesian fine-mapping method to assign causal probability to genes^76,77^.

For FOGS fine-mapping, GTEx V8 Elastic net model based prediction models (weights) were imported from PredictDB Data Repository^74^ for the following 10 GTEx tissues, which appeared to be the most relevant tissues to ageing based on our TWAS results: Subcutaneous adipose tissue, Aortic artery, Brain cerebellar hemisphere, Cultured fibroblast cells, Not sun exposed suprapubic skin, Skeletal muscle, Tibial nerve, Small intestine terminal ileum, Thyroid and Whole blood.

FOGS-type weight and gene lists were generated for each of the 10 GTEx tissues separately^75^. For the LD reference panel, the 1000 Genomes European reference panel was used. Loci definition was from LDetect^78^, where each row in the .bed file represents a single block (start, stop).

TWAS fine-mapping was performed on the parental lifespan^31^ discovery study. The significant (FDR < 0.1) discovery TWAS gene associations were grouped into unique gene clusters based on their chromosomal location using a <100kb window. TWAS-fine mapping was performed on gene clusters with two or more genes, for each of the 10 GTEx tissues separately. The 10 GTEx tissue models used for fine-mapping were chosen as a combination of observing the top 10 highest number of significant TWAS gene associations in the tissues, and having the 10 highest RNASeq and genotyped samples in GTEx V8. The top two genes from each cluster, with the smallest TWAS p-values had to be available in the FOGS fine-mapping results in the same tissue(s) to consider fine-mapping results valid. In gene clusters where this condition was not met, we concluded that the genomic region (using the current input tissue models) cannot be fine-mapped. To determine the putative causal gene in the region, FOCUS PIP scores (generated by FOGS) were analysed, and a gene was considered as putatively causal when the FOCUS PIP score was >0.8 (>80 % probability of being causal). Genes that were located physically distant from any single multi-gene cluster (> +/- 100kb) were considered as independent hits.

### SEEK gene co-expression analysis

SEEK is a gene-based human co-expression search system^79^ containing thousands of expression datasets. SEEK provides co-expression scores (distributed from -3 to +3 for single gene queries) representing z-scores based on rank-biased version of Pearson correlation^79^. Seek gene co-expression analysis was performed on genes of the multi-gene clusters of the parental lifespan discovery TWAS. Genes within a multi-gene cluster were considered significantly co-expressed, if the SEEK co-expression score was ≥ 1.96 (corresponding to 5% probability for a two-sided test) for single gene comparison within a cluster, or across a multi-gene cluster. We also performed SEEK co-expression analysis as a sensitivity analysis for genes considered independent (located in physically distant loci (> +/- 100kb)) and used the same cut-off threshold as above to confirm co-expression status amongst these genes.

### Definition of replicating genes and novel genes

P-values of the parental lifespan TWAS were adjusted using Benjamini & Hochberg false discovery rate (FDR) correction using a 10% cut off threshold. FDR correction was performed in R, using the “p.adjust” function^80^. To identify replicating genes, FDR significant (p<0.1) genes of the discovery TWAS were searched in the healthspan and longevity TWAS. Genes that passed the replication threshold of p<0.05 and had consistent direction of effects (either in the best performing tissue or by the overall mean z-score direction of effect of multiple tissues) in one or both of the other ageing outcomes (i.e. healthspan, longevity) were considered as replicating genes.

To identify novel replicating, ageing-associated genes, the genomic locations of GWAS hits from previous ageing GWAS^13,14,17,20,23^ were queried against the start and end positions (+/-100kb) of the FDR significant (p<0.1) genes of the discovery TWAS. Genes overlapping with the previously identified GWAS loci were considered as established gene associations, while non-overlapping genes were considered as novel gene associations with ageing outcomes tested in this study.

### Gene Ontology analysis

Gene ontology (GO) analyses were performed using Enrichr^32,81,82^, Enrichr-KG, which combines selected gene set libraries from Enrichr for integrated analyses^83^.

We performed GO analysis on the parental lifespan genes replicating in healthspan and longevity. To account for the potential co-expression of nearby genes, we used a 100kb window to group gene associations located near each other, creating gene clusters. The independent genes (based on location); and the strongest gene associations with the smallest p-values from each multi-gene cluster (73/106 genes from healthspan replication and 37/47 genes from longevity replication) were taken forward to the GO analyses.

Parental lifespan genes replicating in healthspan and longevity were analysed separately in Enrichr, using the “Ontologies → Biological Processes” function and were visualised as bar graphs using the clustergram option. All ontology analyses were performed on 09/09/2023.

### Gene selection for the fly experiments

Four of the strongest and replicating human genes, *TOMM40, MAD2L1, COASY and SH3PXD2A* identified through TWAS was selected for the fly experiments based on the following criteria: We first grouped our strongest TWAS associations into two main groups: Group 1: “Parental lifespan genes replicating in both healthspan and longevity” and Group 2: “Novel and highly significant parental lifespan genes replicating in healthspan”. We selected the strongest genes with the smallest p-values from these two groups where genetic tools for *Drosophila* were also available. Our selection was informed by biological relevance and was based on the availability of suitable fly orthologues and genetic reagents.

### *Drosophila melanogaster* orthologue identification

From the list of human genes associated with ageing, we identified four orthologous genes found in *D. melanogaster* using the database Orthologous Matrix (OMA) Browser^84^ (Supplementary Table 10a) which uses pairwise analyses to discover orthology^85^. Based on the direction of effects of gene association obtained from the TWAS analysis, we used RNAi or overexpression lines to modulate the expression of the target genes in the specific tissues in a way that TWAS results predict would increase lifespan. Orthologues with the highest confidence scores were selected and shown in Supplementary Table 10a.

### Expected proportion of lifespan determining genes in model organisms

To estimate the proportion of genes implicated in lifespan regulation, we queried the AmiGO Gene Ontology database^86^ using the term “determination of adult lifespan” (GO:0008340). In *Drosophila melanogaster*, 209 of approximately 13,600 genes were annotated with this term (∼1.54%), while in *C. elegans*, 454 of ∼20,000 protein-coding genes were annotated (∼2.27%). We also included findings from two large-scale RNAi screens: Curran and Ruvkun (2007) identified 64 lifespan-extending genes from 2,700 essential developmental genes (∼2.37%)^44^, and Hansen et al. (2005) identified 23 lifespan-extending genes from a screen of 16,757 genes (∼0.14%)^45^.

### Fly stocks and reagents

Fly stocks were reared and all experiments were performed on sugar/yeast/agar (SYA) medium in a controlled environment set to 25°C, constant humidity of 60% and a 12 h light:12 h dark cycle^87^. The Gene-Switch system was used to induce gene expression in specific tissues^34,35^. The Gene-Switch drivers used to drive expression ubiquitously, in the gut, and neurons in this study are daughterless-GS (da-GS), TIGS, and elav-GS respectively. All driver stocks used were backcrossed at least 6 times into the healthy, outbred Dahomey background carrying a *w*^1118^ mutation and are *Wolbachia* negative. RNAi and overexpression lines were obtained from the Bloomington Stock Centre^88^. RNAi lines used include *Ppat-dpck* (FBgn0035632), *Tom40* (FBgn0035632), *cindr* (FBgn0027598), and a control RNAi targeting *RFP* (FBgn0044485), while *mad2* (FBgn0033809) overexpression line was used. These were not backcrossed, except the RNAi line targeting *Ppat-dpck* in the second experiment, which was backcrossed 6 times into the outbred Dahomey background carrying a *v^1^* mutation to allow the tracking of the transgene. To validate the RNAi or over-expression constructs, we used *daughterless-GAL4*. To generate the experimental flies used in this study, the male flies from the RNAi and overexpression lines were crossed with the females of the Gene-Switch drivers. Genotypes of the lines used along with their Bloomington stock numbers are available in Supplementary Table 10a.

### *Drosophila melanogaster* lifespan assay and analysis

Experimental flies were reared at standard larval density. Emerged adult flies were left to mate for 48 hours before being sorted by sex under brief CO_2_ anaesthesia and transferred to experimental vials (on SYA medium containing zero or 200 µM RU486) at a density of 15 female flies per vial. The flies were transferred to vials containing new food every 2 – 3 days and the age and deaths were recorded until the last survivor was dead. Lifespan data were subjected to survival analysis by log-rank test and presented as survival curves.

### Quantitative PCR

Mothers carrying the ubiquitous, constitutive driver *daughterless-GAL4* were crossed to the males carrying the RNAi or overexpression constructs, or a control not carrying any transgenic construct. RNA was extracted with Trizol (Invitrogen) from larvae (for Mad2, Tom40 or Cindr) or young adults (Ppat-dpck) and converted to cDNA using oligo-dT and Superscript II (Invitrogen) according to manufecturers’ instructions. Quantitative PCR was performed using Power SYBR Green PCR Master Mix (ABI), on QuantStudio6 Flex real-time PCR, with the relative standard curve method. Primers used were: *Mad2_*Forward (ACCTGAAGTACGGCATCAATTC) and *Mad2*_Reverse (GTTTGGCTGAGCACGTTCTG), *Cindr_*Forward (GCGGAGTTCCTATGACTGGTA) and *Cindr*_Reverse (GCAAGACCATCCGCCAAA), *Tom40*_Forward (CCCTCTAATTCCGGCTCACT) and *Tom40*_Reverse (TGGCCTGGATATCTTTGC), *Ppat-dpck*_Forward (GCCATGACGAAGGGAAAGAC) and *Ppat-dpck*_Reverse (GGAACATTCGATCGCATCCA), *Actin*_Forward (CACACCAAATCTTACAAAATGTGTGA) and *Actin*_Reverse (AATCCGGCCTTGCACATG).

## Data availability

We used publicly available data for the analyses. Summary statistics for the parental lifespan GWAS, healthspan GWAS and longevity GWAS are available at https://www.ebi.ac.uk/gwas/home under study accession IDs GCST009890, GCST007406, GCST008599, respectively. All data obtained in fruit fly experiments are available in the supplement.

## Code availability

We used publicly available software for the analyses. The software used is listed in the Methods section.

## Acknowledgements

We are grateful to all the participants who took part in the studies included in this work. We would like to thank all members of the UCL HumGen lab (https://www.uclhumgen.com/), who gave critical support and suggestions. This work was conducted on the University College London Computer Science cluster, we would like to thank the cluster team for the support provided. Stocks obtained from the Bloomington Drosophila Stock Center (NIH P40OD018537) were used in this study and the RNAi lines were made by the TRiP project (Office of the Director R24 OD030002: “TRiP resources for modeling human disease” PI: N. Perrimon).

## Author contributions

G.N., K.K. and N.A. conceived this project. K.K., N.A., and L.P. supervised the work. G.N. wrote the manuscript. G.N., K.K. and N.A. critically revised the manuscript. G.N. carried out the TWAS, Fine-mapping, Gene co-expression and Gene Ontology analyses. A.A. carried out the experimental work in *Drosophila melanogaster.* O.G. and S.M. provided support for the TWAS. All authors read and approved the manuscript.

